# Selection removes Shine-Dalgarno-like sequences from within protein coding genes

**DOI:** 10.1101/278689

**Authors:** Adam J. Hockenberry, Luίs AN Amaral, Michael C. Jewett, Claus O. Wilke

## Abstract

The Shine-Dalgarno (SD) sequence motif facilitates translation initiation and is frequently found upstream of bacterial start codons. However, thousands of instances of this motif occur throughout the middle of protein coding genes in a typical bacterial genome. Here, we use comparative evolutionary analysis to test whether SD sequences located within genes are functionally constrained. We measure the conservation of SD sequences across Gammaproteobacteria, and find that they are significantly less conserved than expected. Further, the strongest SD sequences are the least conserved whereas we find evidence of conservation for the weakest possible SD sequences given amino acid constraints. Our findings indicate that most SD sequences within genes are likely to be deleterious and removed via selection. To illustrate the origin of these deleterious costs, we show that ATG start codons are significantly depleted downstream of SD sequences within genes, highlighting the potential for these sequences to promote erroneous translation initiation.

## Introduction

The Shine-Dalgarno (SD) sequence is a short motif that facilitates translation initiation via direct base pairing with the anti-Shine-Dalgarno (aSD) sequence on the 16S ribosomal RNA [1]. Several previous studies have shown that SD sequences are significantly depleted from *within* the protein coding genes of many bacterial species [2–4]. Although the depletion of SD sequences within protein coding genes is highly *statistically* significant, many prokaryotic genomes nevertheless contain thousands of these sequences While SD sequences and their effect on translation initiation have been studied for decades [6–8], the role of these SD sequences within protein coding genes—hereafter referred to as SD-like sequences—is relatively unknown.

SD-like sequences may promote spurious internal translation initiation resulting in the production of truncated or frame-shifted protein products that are likely to be deleterious [9]. These sequences are also known to promote ribosomal frame-shifting during translation elongation, which can have a beneficial regulatory function in specific cases [10–12]. More recently, researchers have suggested a general role for SD-like sequences in regulating the rate of translation elongation [13]. The evidence for translational pausing at SD-like sequences is supported by ribosome profiling studies in several bacterial species [13–16] as well as experimental studies using a variety of techniques If SD-like sequences regulate elongation rates, many of the observed SD-like sequences within genes may actually be beneficial for cells; translational slowdown and pausing has been shown to facilitate proper protein folding in a number of different contexts [22–30].

However, other researchers have hypothesized that the experimental evidence for an association between SD-like sequences and translational pausing in ribosome profiling data may be an experimental artifact rather than a true biological effect Using a variety of different experimental techniques, other studies have failed to observe an association between the appearance of SD-like sequences and ribosomal pausing events [32–37].

Taken together, the experimental evidence for whether SD-like sequences regulate translation elongation rates is mixed. Further, if this mechanism of translational pausing is real, we still would not know whether organisms rely on the presence of SD-like sequences to regulate the rate of translation elongation. Just as plausibly, the cellular costs related to frame shifting and spurious initiation may outweigh any benefits that would arise from employing this regulatory strategy. Determining the balance of these various effects is important for recombinant protein production applications that could use knowledge of SD-like sequences to tune elongation rates and encourage the production of properly folded proteins

Here, we apply comparative evolutionary analysis to determine whether SD-like sequences in the genome of *E. coli* are deleterious, neutral, or beneficial. Evidence for conservation of these sequences would indicate that they are beneficial, perhaps due to a regulatory role in translation elongation. By contrast, our results show that 4-fold redundant codons within SD-like sequences have significantly *higher* substitution rates than expected according to two different null model controls. These findings hold across a number of attempts to isolate a pool of functionally constrained sites, and strongly suggest that SD-like sequences are weakly deleterious throughout the *E. coli* genome. We find that start codons are significantly depleted downstream of existing SD-like sequences, which provides evidence for the deleterious effects related to internal translation initiation that these sequences may promote. Our findings cast doubt on the role of SD-like sequences as a potential regulator of translation elongation rates in native genes, and urge caution when employing methods that use these sequences to tune translation elongation in recombinant designs.

## Results

### Assessing the conservation status of Shine-Dalgarno-like sequence motifs within protein coding genes

To investigate whether SD-like sequence motifs that occur within protein coding genes have a functional role, we searched for signatures of evolutionary conservation of these sites across related species. Under the hypothesis that some fraction of the SD-like sequence motifs that are present in any genome may be playing an important functional role, we would expect to observe significantly lower rates of nucleotide substitution within these sequence motifs relative to control sites. Conversely, if these sequences perform no such functional role and are instead generally deleterious to organismal fitness, we should observe significantly higher rates of substitution within these sequence motifs.

We assembled a dataset of 1394 homologous protein families from 61 species in the order *Enterobacterales* and quantified nucleotide-level substitution rates across the coding sequences from this dataset. We used *E. coli* as a reference organism to identify the location of all SD-like sequence motifs that contain 4-fold redundant nucleotide sites in conserved amino acid positions while ignoring sites at the 5ʹ and 3ʹ gene ends (see Materials and Methods). We note that canonical SD sequences are often not perfect complements to the highly conserved anti-SD sequence [38–40], and in this manuscript, unless specified otherwise, we used a binding energy threshold of 4.5 kcal/mol to define SD-like sequences. According to this threshold, 1998 out of 4127 *E. coli* protein coding genes are preceded by SD sequences, significantly more than expected by chance (Expectation: 638.57, *z*-test:*p <* 10 ^16^). By the same definition, all *E. coli* protein coding genes contain 25,001 SD-like sequences, significantly fewer than expected by chance alone (Expectation: 30,397.57, *z*-test:*p <* 10 ^16^) but far more than the number of known SD sequences that function in translation initiation (Supplementary Table S1).

We adopted a paired-control strategy to compare substitution rates between nucleotide sites that fall within SD-like sequence motifs to control sites selected from the same gene that *do not* occur within SD-like sequences. Throughout the remainder of this manuscript we use the nomenclature of ‘codon’ and ‘context’ controls to refer to two different methods for selecting control nucleotides. In codon controls, after identifying a 4-fold redundant codon *within* a SD-like sequence, we find another occurrence of the same codon within the same gene to use as a control. Similarly, in context controls we find the same tri-nucleotide site (at the -1, 0, and +1 positions, where a 4-fold redundant position is at position 0) within the same gene to use as a paired control (Fig. 1A). These two null models control for possible effects arising from synonymous codon usage bias and biases that may emerge from mutational context, respectively.

**Figure 1:**
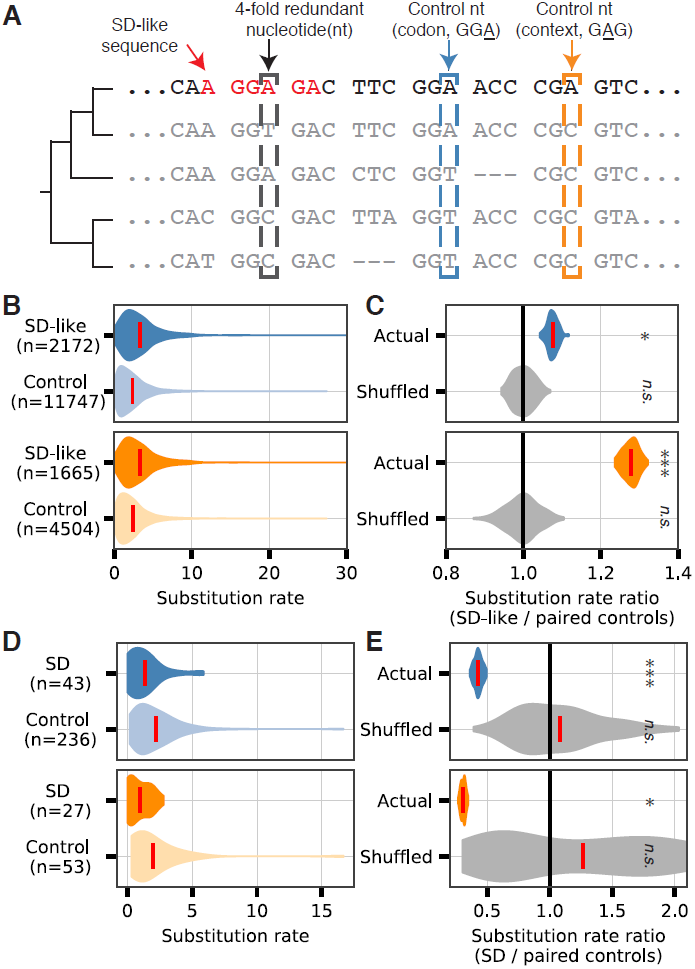
SD-like sequences have elevated rates of nucleotide substitution. **(A)** Graphical illustration of methodology for identifying 4-fold redundant sites within SD-like and control sites. **(B)** Relative substitution rates for all SD-like sites and control sites. Top (blue) and bottom (orange) panels depict results for codon and context controls, respectively. Red lines in violin plots depict category means. **(C)** The ratio of the average substitution rates between SD-like and control categories based on a gene-specific bootstrapped approach discussed in main text (*p* = 0.003, 0.48, 1.2 10 ^-14^, 0.53, top to bottom). **(D)** As in (B), showing scores for putative SD sites within the 3 ^ʹ^ end of genes. **(E)** Substitution rate ratios for putative SD sites depicted in (D)(p = 0:001; 0:48; 0:013; 0:51, top to bottom). (*denotes p < 0:05, ***denotes p < 0:001)

Since SD-like sequences are relatively rare, there are frequently many possible control sites within a given gene for each synonymous nucleotide that occurs within a SD-like sequence (Fig. 1B). We thus randomly sampled single control nucleotide sites (from within the same gene) for each applicable SD-like nucleotide. From the resulting paired list of substitution rates for SD-like and control sites, we calculated the ratio of the average substitution rates between the two categories (SD-like sites divided by control sites) and repeated this sampling procedure 100 times to estimate the overall effect size. Assuming no difference in substitution rates between SD-like and control sequence categories, the ratios should follow a normal distribution centered around a value of 1. If sites within SD-like sequences are more conserved than control sites, we should observe values significantly less than 1. Finally, if sites within SD-like sequences have elevated substitution rates, indicating that they are generally deleterious, we expect to observe ratios significantly greater than 1.

Regardless of which null model strategy that we used to select control nucleotides, we found that the substitution rates of SD-like sequences are *higher* than that of control sequences with an effect size on the order of ~10–30% (Fig. 1C). By contrast, when we randomly assigned nucleotide sites to SD-like or control categories the resulting distribution of substitution rate ratios was centered around the expected value of 1 (‘shuffled’ data in Fig. 1C). We conservatively determined statistical significance by calculating the Wilcoxon signed-rank test between SD-like and control categories for each bootstrap replicate and report the median *p*-value (*p* = 0.003 and *p* = 1.2 ×10 ^-14^*-*^14^ for codon and context controls, respectively). These results remain qualitatively unchanged when we used different thresholds to define SD-like sequences (Supplementary Fig. S1), as well as a different organism (*Y. pestis*) to identify the locations of SD-like sequences (Supplementary Fig. S2).

To ensure that our methodology was capable of predicting conservation of sequence motifs that are *known* to be functionally constrained, we leveraged the fact that some genes in our dataset are directly followed by another gene in the 3ʹdirection. Thus, the SD sites of certain downstream genes are expected to occur within the 3ʹ coding sequence of upstream genes. We therefore repeated our analysis by considering only putative SD sites that occur within the 50 to 1 region (relative to the stop codon) in the subset of genes where another gene directly follows (while still selecting control sites from the internal regions of the gene). Despite the low number of motifs that met this criterion, 4-fold redundant sites within this restricted set of putative SD sequences had a substitution rate that is roughly 1*/*3 that of control nucleotides, indicating strong evolutionary conservation of these known SD sites and validating our overall statistical approach (Fig. 1D,E). We ensured that this result was not simply an artifact of differential substitution rates at the 3ʹ end of genes by conducting the same analysis on sites that occur within the 3ʹ region of genes that *do not* have any annotated genes directly following, and thus are not expected to function as true SD sites. We detected no significant signal of evolutionary conservation for this set of sites (SI Fig. S3).

### Substitution rates differ according to mutational effects on SD-like sequence strength

In the preceding section, we showed that 4-fold redundant sites within SD-like sequences have significantly higher substitution rates than control sites. This finding provides support for the model of SD-like sequences being deleterious and evolutionarily transient within genes. However, the SD sequence binds facilitates translation initiation by binding directly to the anti-SD(aSD) sequence on the 30S ribosomal subunit, and this binding strength spans a range of values according to the actual SD nucleotide sequence in question. We thus separately investigated SD-like sites according to how many synonymous mutations to the 4-fold redundant nucleotide in question were predicted to increase the strength of binding to the aSD sequence (see Fig. 2A for an example). Note that this designation does not refer to the absolute strength of aSD sequence binding, but rather the capacity for *strictly* synonymous mutations to the site in question to either increase or decrease the relative aSD sequence binding. We refer to the ‘locally strong’ and ‘locally weak’ sites hereafter as those where any synonymous mutation is guaranteed to decrease or increase, respectively, the strength of aSD sequence binding.

**Figure 2:**
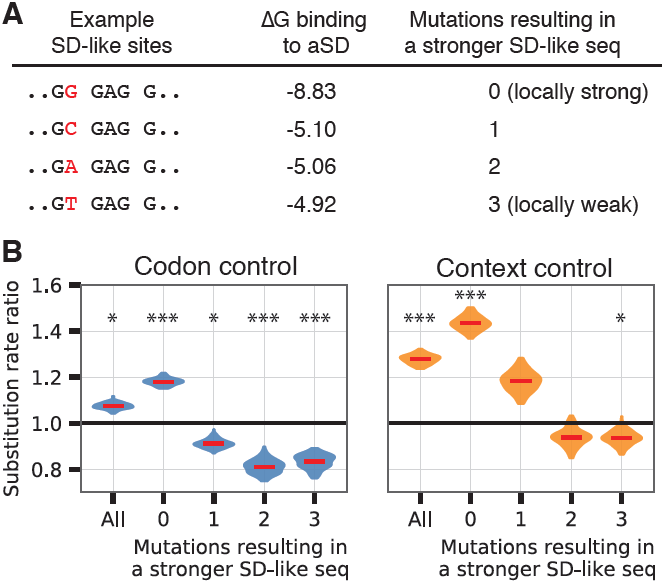
Local mutational effects on SD strength alter substitution rate patterns. **(A)** Synonymous mutations to SD-like sequences may either increase or decrease SD-like sequence strength depending on the identity of the 4-fold redundant nucleotide. **(B)** Substitution rate ratio results as in Fig.1C. Data shown here by stratifying ‘all’ sites into categories that correspond to the expected change in SD strength given a synonymous substitution. Results shown for synonymous codon (left, (*p* = 0.003, 2.7 ×10 ^-12^, 0.006, 0.0004, 4.2×10 ^-5^) and nucleotide context (right, *p* = 1.2×10 ^-14^, 1.3×10 ^-22^, 0.17, 0.2, 0.026) controls.(* denotes *p <* 0.05, ***denotes *p <* 0.001)

Based on our previous results, we hypothesized that if SD-like sites are deleterious, we should observe conservation of locally weak sites. For this subset of sites, any synonymous mutation would, by definition, result in an *increased* aSD sequence binding strength. Indeed, substitution rates for this category of sites were significantly lower than expected (substitution rate ratios less than 1, *p <* 0.01), regardless of our method for selecting control nucleotides (Fig. 2B). By contrast, when we analyzed the subset of locally strong SD-like sites, where any mutation to the 4-fold redundant position is guaranteed to result in a *weaker* interaction with the aSD sequence, we observed the opposite effect. These sites—which are the majority of identified SD-like sites—had substantially elevated substitution rates compared to paired controls on the order of ~10–40% (see Table 1 for the number of data points included in each category, which are highly skewed towards locally strong sites in this analysis).

We stress that these findings are not indicative of conservation of intermediate or weak SD-like sites, but rather the *weakest possible* sites given the amino acid constraints of the sequence. To further address this point, we performed the same analysis on weak SD-like sites, which we define as having aSD sequence binding free energy values between 3.5 and 4.5 kcal/mol. We observed the same pattern of locally strong sites having significantly elevated substitution rates; this is despite the fact that these sites are weaker in absolute terms than all sites depicted in Fig. 2 (SI Fig. S4). This nucleotide dependent analysis shows that the magnitude of negative selection acting against SD-like sites is stronger than we initially observed in Fig. 1. As before, to ensure the robustness of these results we we used different thresholds to define SD-like sequences (Supplementary Fig. S5), as well as a different organism (*Y. pestis*) to identify the locations of SD-like sequences and their classifications (Supplementary Fig. S6) and observed consistent results.

### Consistent results across protein abundance bins

While we have thus far shown that SD-like sequences as a whole are less conserved than expected, this does not preclude the possibility that some fraction of SD-like sequences have a functional role and are evolutionary constrained. The SD-like sequences that we have analyzed may actually be a mixture of deleterious and functionally beneficial sites that look weakly deleterious in aggregate. We reasoned that the most highly abundant proteins are most likely to have been purged of deleterious SD-like sequences leaving the SD-like sequences that remain within these genes particularly attractive candidates for functional conservation. Thus, if SD-like sequences are a mixture of effects, we expect to find SD-like sites within highly expressed genes to be *relatively* more conserved than other categories. By contrast, if SD-like sites are a uniform pool in terms of their overall negative effects, we predict that the substitution rates between different gene expression categories will not systematically vary. To test this hypothesis, we separated our dataset into quintiles of genes according to their overall protein abundances in *E. coli*, and analyzed the substitution rate ratios of SD-like and control categories as before.

We confirmed that the most highly abundant proteins contain fewer SD-like sequences (Fig. 3A). Since the the fraction of conserved amino acids per gene varies according to bins of protein abundance (Fig. 3B), the overall fraction of SD-like sites eligible for analysis is variable between different protein abundance bins (Fig. 3C). However, we nevertheless observed largely consistent results across all protein abundance bins: locally strong 4-fold redundant nucleotides within SD-like sequences have significantly higher substitution rates than paired controls (Fig. 3). These results remained robust to our assumptions with regard to SD-like thresholds (Supplementary Fig. S7) and species used to identify SD-like sites (Supplementary Fig. S8)—though we note in the latter case *E. coli* values were still used to classify homologs into protein abundance bins. We also found that the locally weak SD-like sites had significantly lower substitution rates than expected across nearly all protein abundance bins with the only exceptions being for the sites within the very lowest protein abundance bins (SI Fig. S9).

**Figure 3:**
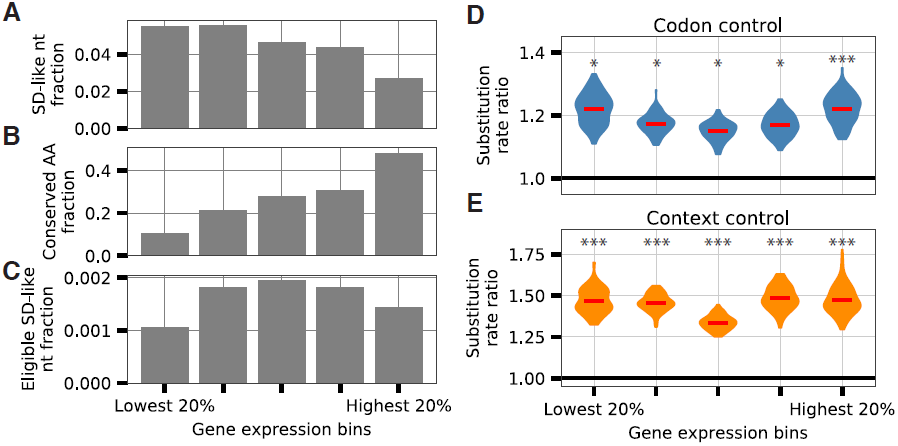
SD-like sequences have similarly elevated substitution rates across protein abundance bins. **(A)** The most highly abundant proteins contain fewer SD-like sequences. **(B)** Highly abundant proteins are have a higher fraction of conserved amino acids. **(C)** Combined, the effects from (A) and (B) affect the fraction of SD-like sites within genes that are eligible for our analysis. **(D)** Substitution rate ratios of the locally strong SD-like sequences are elevated across all levels of protein abundance compared to synonymous codon controls (*p* = 0.009, 0.003, 0.004, 0.001, 0.0005). **(E)** As in (D), shown according to context controls(*p* = 0.0001, 2.3×10^-6^, 1.9×10^-5^, 8.4×10 ^-7^, 4.0×10 ^-5^). (* denotes *p<*0.05, ***denotes *p <* 0.001) is highly similar, casting doubt on the hypothesis that SD-like sites within a genome are actually composed of a mixture of functionally constrained and deleterious sites.

Importantly for our goal of trying to delineate between competing hypotheses, we found no evidence of a consistent trend that would indicate that sites within highly expressed proteins were more or less likely to show evidence of functional constraint. By contrast, the overall pattern of relative substitution rate ratios across different protein abundance bins

**Table 1.**
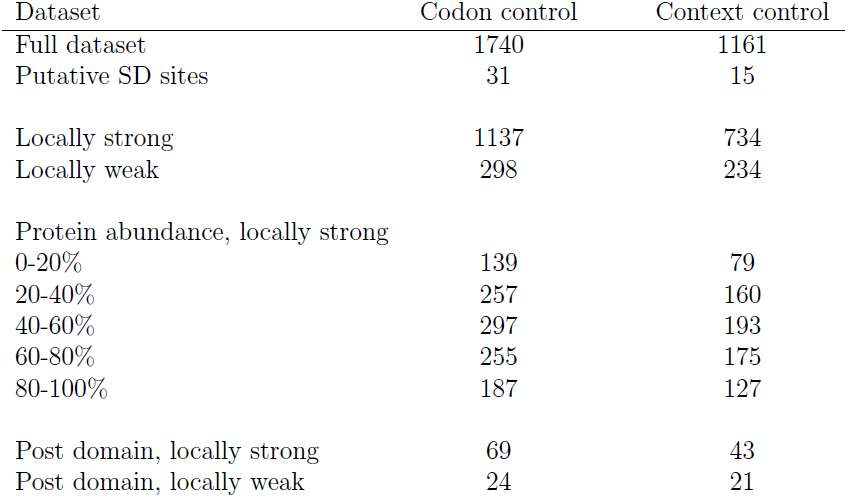
Number of sites analyzed in each bootstrap replicate used to calculate substitution rate ratios. Note that this number differs from the total number of all SD-like sites identified for a given criteria as some SD-like sites lack suitable control sites and are discarded from further analysis. Additionally, when the number of SD-like sites in a gene exceeds the number of control sites for a given criteria, pairs of sites are randomly sampled without replacement until no control sites remain for a given bootstrap replicate and all further SD-like sites discarded from analysis for that particular replicate.

| Dataset | Codon control | Context control |
| --- | --- | --- |
| Full dataset | 1740 | 1161 |
| Putative SD sites | 31 | 15 |
| Locally strong | 1137 | 734 |
| Locally weak | 298 | 234 |
| Protein abundance, locally strong |  |  |
| 0-20% | 139 | 79 |
| 20-40% | 257 | 160 |
| 40-60% | 297 | 193 |
| 60-80% | 255 | 175 |
| 80-100% | 187 | 127 |
| Post domain, locally strong | 69 | 43 |
| Post domain, locally weak | 24 | 21 |

### Consistent results for sites following protein domain boundaries

Most studies that have explored the possible functional benefits resulting from elongation rate variability have focused on the role that slow translation or translational pausing may have in helping to enhance co-translational protein folding. Past research has indicated that slow translation at domain boundaries may enhance protein solubility by allowing one domain to properly fold before the next domain fully emerges from the ribosome exit tunnel The most probable candidates for functional SD-like sites may thus be those sites that occur after protein domains.

To test this hypothesis, we relied on previously curated protein domain annotations from Ciryam *et al.* (2013)[41]. After merging datasets, we were left with 415 proteins in our dataset with domain annotations. We repeated our analysis within this subset of proteins, while only considering SD-like sites that occur after protein domains. We define this region as the 30-150 nucleotides downstream of 3ʹ domain boundaries to account for uncertainty in annotations, and maintained our previous restriction of discarding data from the first 100 and the last 50 nucleotides for each gene. We specifically looked at the locally strong and locally weak sites, expecting that these categories would show the strongest signal based on our findings in Fig. 2B.

Under the hypothesis that SD-like sites after protein domains may have a functional role, we expected to observe conservation of this subset of SD-like sites (substitution rate ratios less than 1). A slightly weaker version of this hypothesis is that these SD-like sites should be *relatively* more conserved than SD-like sites overall. If instead SD-like sites following protein domain boundaries do not represent any special category of sites, we should observe results similar to our prior findings where we observed elevated substitution rates in locally strong sites and conservation of locally weak sites.

For both codon and context controls, we found that substitution rates are significantly greater than 1 for locally strong sites following protein domains with no substantial difference between these sites and the aggregated set of all locally strong SD-like sites (Fig. 4A). Our results for locally weak sites were also consistent with the hypothesis that SD-like sites following protein domains are not obviously a distinct category of SD-like sites (Fig. 4B). In both cases, we found more heterogeneity in the estimates for the mean substitution rate ratios for the post-domain categories, and note that this reflects the comparably small number of SD-like sites that meet the relevant criteria for this analysis (Table 1).

**Figure 4:**
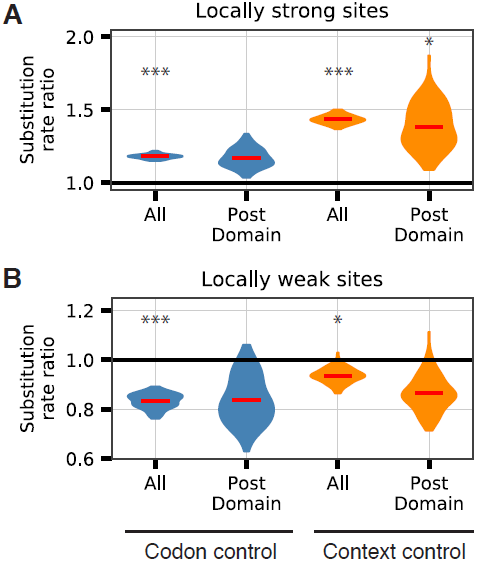
Consistent results following protein domain boundaries. **(A)** Locally strong SD-like sites downstream of protein domain boundaries exhibit elevated substitution rates similar to all SD-like sites (*p* = 2.7×10 ^-12^, 0.2, 1.3×10 ^-22^, 0.04). **(B)** Similar results to (A) for locally weak SD-like sites following protein domain boundaries (*p* = 4.2×10 ^-5^, 0.4, 0.026, 0.3). The greater heterogeneity for post-domain sites in both panels reflects the comparably small number of sites meeting the indicated criteria (see Table 1). (* denotes *p <* 0.05, ***denotes *p <* 0.001)

**Figure 5:**
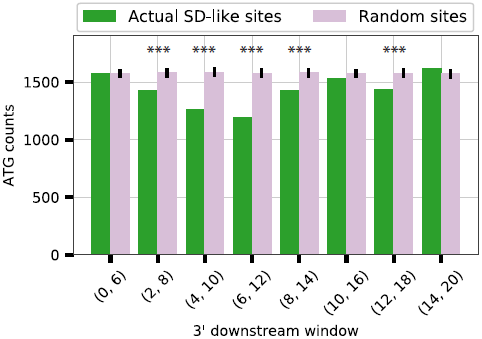
Start codons are depleted downstream of SD-like sequences. We tallied the number of ATG tri-nucleotide sequences that occurred within the indicated windows downstream of SD-like sequences throughout the *E. coli* genome. An equivalent number of random sites within each gene were selected as a control calculate significance (*p* = 8.6 ×10^-5^, 3.6 ×10 ^-15^, 5.2 ×10^-27^, 9.3 ×10^-5^, 0.0003 for comparisons marked as significant). (***denotes *p <* 0.001)

### SD-like sequences and internal translation initiation

All of our results with regard to sequence conservation point to SD-like sequences having elevated rates of substitution indicative of their being largely detrimental to long-term cellular fitness. But exactly what are these detrimental effects? A natural hypothesis is that SD-like sequences may result in erroneous translation initiation, which would produce truncated or frame-shifted protein products. To test whether there is evidence of this effect, we extracted nucleotide sequences downstream of all SD-like sites within the *E. coli* genome (*n* = 25, 001). For a given downstream window, we asked how many ATG tri-nucleotide sequences occur (regardless of reading frame). We observed a significant depletion of ATG tri-nucleotide sequences within a relatively narrow window downstream of SD-like sites (4-12 nucleotides) that is in line with expectations from the characteristic spacing observed in true SD sites [43]. We calculated random expectation by drawing an equivalent number of random locations per-gene, performing the same analysis, and repeating this procedure 100 times. We observed no qualitative decrease in ATG counts according to this null model at different windows and calculated the significance of each window in the observed data according to this null expectation using a *z*-test. These results show that coding sequence patterns are constrained as a result of SD-like sequence occurrence so as to minimize possible translation initiation events. The detrimental effects of such erroneous translation initiation events likely explain at least part of the selection against the occurrence of SD-like sequences within protein coding genes.

## Discussion

Several previous studies have shown that SD-like sequences are somewhat depleted within the protein coding genes of bacteria These studies, however, could not comment on whether SD-like sequences are deleterious to organismal fitness or whether they are sparingly used because they serve a potentially important regulatory function. Recently, there has been a debate in the literature as to the possible role that SD-like sequences may play in regulating translation elongation rates with different experimental protocols yielding conflicting results [13, 32]. Here, we pursued a complementary approach to investigate the possible function of SD-like sequences within bacterial protein coding genes. We performed a comparative evolutionary analysis and found that SD-like sequences are weakly deleterious throughout the *E. coli* genome.

Using a relatively strong definition to classify SD sequences, we found that roughly 2,000 of the 4,000 *E. coli* protein coding genes are preceded by an identifiable SD sequence slighty upstream of the start codon (Table S1). This is substantially more than the ~600 that would be expected based off the nucleotide composition of UTRs. However, according to this same definition, there are nearly 25,000 SD-like sequences scattered throughout *E. coli* protein coding genes (after excluding the first and last 60 nucleotides). The number of these SD-like sequences is significantly fewer than the ~30,000 that would be expected based off of codon usage biases and amino acid sequences, but the overall magnitude of depletion is relatively modest in scale. While these exact numbers are subject to change based on various thresholds and definitions, the facts remain that (i) there are greater than 10 times more SD-like sequences inside *E. coli* protein coding genes than there are true SD sequences, (ii) the overall depletion of SD-like sequences relative to expectation is highly significant yet small in magnitude, and (iii) in the majority of cases, we do not know whether the existing SD-like sequences have any function at all.

Sequence conservation remains one of the gold standards for assessing the functionality of DNA sequences or regions [44–46]. We therefore looked at the evolutionary conservation of 4-fold redundant sites that occur within SD-like sequences across *E. coli* protein coding genes. We compared the conservation sites within SD-like sequences to gene-specific control sites to determine whether there was any evidence of functional constraint. We failed to find any evidence of evolutionary conservation for the set of all SD-like sequences within our dataset of Gammaproteobacteria, and instead found that these sequences actually have significantly elevated rates of substitution, on the order of ~10–40% depending on the method used to select control sites. In addition to looking at all SD-like sequences, we performed a number of robustness checks and attempted to isolate subsets of likely functionally constrained SD-like sequences. However, considering sets of SD-like sequences according to (i) their overall strength of binding to the aSD sequence, (ii) their occurrence within highly or lowly expressed genes, or (iii) their locations relative to known protein domain boundaries did not alter our findings.

By contrast, we know that some SD-like sequences are functional and we did find that SD-like sequences that are *true* SD sequences for downstream genes in multi-gene operons are highly conserved. We also found that *locally weak* SD-like sequences are conserved; in these sequences, any mutation to the 4-fold redundant site in question would actually result in an increased SD-like strength. Conservation of nucleotides within these locally weak sites is therefore evidence for avoidance of strong SD-like sequences and supports our conclusion that SD-like sequences are generally deleterious.

Researchers have previously shown that SD-like sequences are capable of promoting internal translation initiation [9]. We therefore hypothesized that the deleterious effects of SD-like sequences may be due to their role in encouraging internal translation initiation which would create truncated and/or frame-shifted protein products. Indeed, we found strong support for this hypothesis by observing that the occurrence of ATG start codons is significantly depleted within a narrow window downstream of existing SD-like sequences in *E. coli*. These data suggest that when SD-like sequences appear, they induce additional downstream constraints on coding sequence evolution and these constraints are consistent with the avoidance of translation initiation sequence features.

Since our analyses were performed on aggregates of SD-like sequences, we could not rule out whether any individual SD-like sequence or any particular set of sequences are highly conserved. In fact, we observed numerous examples of 4-fold redundant sites within SD-like sequences that are entirely conserved across all 61 species. However, the number of these sites is simply no more (and in fact, substantially fewer) than our two different null model controls. Our results do not rule out the possibility that some alternative grouping of particular genes or regions within genes that we did not consider may show increased conservation compared to null expectation. Nevertheless, based on our results and previously identified examples, the numbers of functionally constrained SD-like sequences that are involved in known regulatory processes—such as programed frame-shifting [10–12]—appear to be a small minority of all the existing SD-like sequences.

While SD-like sequences may cause spurious internal translation initiation, another possible role they may play is in regulating translational pausing [13, 32]. Many studies have argued that pausing during translation can be beneficial, because it may facilitate proper protein folding However, our results here show that the majority of SD-like sequences are likely deleterious. Therefore, we think it is unlikely that SD-like sequences are commonly used as a means to regulate translation elongation and protein folding in endogenous genes. We cannot, of course, rule out that this effect may exist in a limited number of cases.

Taken together, our findings show that SD-like sequences tend to be either purged from closely-related genomes or maintained in their weakest possible state given amino acid sequence constraints. The appearance of so many SD-like sequences throughout bacterial genomes is likely explained by a combination of constant mutational supply, amino acid constraints, and relatively weak selective pressures acting to remove these sequences. Practically speaking, our findings suggest that SD-like sequences should be avoided in the design of recombinant protein expression applications until more is known about their possible deleterious effects to cellular fitness.

## Materials and Methods

### Dataset compilation

We assembled a dataset of 1394 homologous proteins from 61 genomes within the order *Enterobacterales*, unique at the individual species level (see Supplementary Tables S2 for a complete list of analyzed genomes). We chose this set of species as a balance between identifying relatively large numbers of homologous proteins for comparative analysis (which becomes progressively more difficult with more highly-diverged species) while minimizing the confounding effects of population-level polymorphisms that may occur when analyzing multiple members of a single species. We selected species based off of their inclusion in either the PATRIC ‘reference’ or ‘representative’ species designations [52] and used PATRIC-derived gene annotations since these annotations derive from a consistent pipeline. For each genome, we extracted all amino acid sequences and performed a reciprocal USEARCH [53]comparison against *E. coli* amino acid sequences to find 1:1 best hits (using a 70% identity threshold and a strict e-value cutoff of 10 ^-10^). We included all homologs that appeared in at least 45 species.

We next individually aligned the amino acid sequences of each homolog family using MUSCLE [54] and used RAxML (GTR model, 100 bootstrap and 20 maximum likelihood replicates) [55] to create a phylogenetic tree on the concatenated amino acid sequences of 108 genes identified in all species with the fewest number of insertions/deletions. With this tree topology, we next calculated relative nucleotide substitution rates at each position by back translating aligned amino acid sequences into codon sequences and running HyPhy under a GTR model to estimate position-specific substitution rates within each gene. We trimmed any 5ʹ and 3ʹ extensions based on the *E. coli* reference sequence annotations and then normalized each nucleotide substitution rate according to the mean of each gene.

We confirmed that the overall accuracy of relative substitution rate scores by performing several tests. We show via a meta-gene analysis that median substitution rates at 3rd positions of codons are significantly higher than 1st or 2nd positions and that substitution rates at the 5ʹ end of genes are lower than internal positions reflecting selection on mRNA structure surrounding the start codon (SI Fig. S10).

### Quantifying substitution rate differences between motifs

To assess the conservation status of longer sequence motifs while controlling for gene-specific effects, we focused on 4-fold redundant codon sites. We identified SD-like sites according to the computationally predicted hybridization energies between all sequential 6 nucleotide motifs within each gene and a putative anti-Shine-Dalgarno sequence (5ʹ-CCUCCU-3ʹ) using the ViennaRNA [58] co-fold method with default parameters. We used a threshold of 4.5 kcal/mol based-off of the distributions of true SD sequences in the *E. coli* genome (SI Fig. S11) to classify sequences as SD-like.

For each SD-like sequence motif that we identified, we assessed whether there are any 4-fold redundant nucleotide sites present within that sub-sequence (excluding the terminal nucleotides). If so, and if the amino acid site was almost entirely conserved (allowing for one possible amino acid change across the species set) we next found all occurrences of the same synonymous codon within the same gene (so long as it too does not occur within a SD-like motif) subject to the same conservation constraint. We use these 3rd position nucleotides as controls. For both categories (SD-like and matched controls) we excluded nucleotides from our analysis if they fell within 100 nucleotides downstream from the *E. coli* annotated start codon or 50 nucleotides upstream from the stop codon to avoid potentially confounding effects related to translation or termination.

Additionally, we conducted a separate analysis that relied on nucleotide context for selecting control nucleotides. After finding a 4-fold redundant codon in a conserved amino acid site within a SD-like sequence motif as before, we searched for another occurrence within the same gene where there is a 4-fold redundant site with the same nucleotide identity and having the same flanking nucleotides at both the +1 and 1 positions, regardless of whether the synonymous codon is the same (i.e. the 2 position). The rest of the calculation proceeded as above, with the exception that we introduced a further constraint here by requiring the +1 nucleotide to be almost perfectly conserved (less than one substitution) in addition to the amino acid under investigation.

To conservatively estimate the effect size and assess statistical significance between SD-like nucleotides and controls (given their non-normal distribution and unequal *n’*s), we adopted a paired approach as described in the text. For each gene we randomly selected one of the SD-like nucleotide values and one paired-control value (without replacement) until there are either no more SD-like nucleotides or no suitable control nucleotides for the given gene. We then repeated this procedure across all genes in the dataset. This paired analysis method controls for gene-specific effects and creates equally sized categories, which allowed us to estimate the effect size as the ratio between the average relative substitution rates for the SD-like and control site categories. We repeated this sampling procedure 100 times to get a distribution of these ratios and assessed the significance of each bootstrap by performing a Wilcoxon signed-rank test, reporting the median observed *p*-value across all replicates.

Further analyses described in text were performed following the same basic procedure as above, by either stratifying all SD-like sites into categories based on their local mutational effects, their positions within genes, or by classifying sites separately according to different gene sets.

## Protein abundance data

We downloaded protein abundance measurements from the PaxDB database (integrated dataset, accessed 07/2017) [59] and matched gene ids to the PATRIC genome annotation of *E. coli*. We were able to unambiguously map 1,386 of the 1,394 coding sequences in our complete dataset to protein abundance measurements. We split these into equally sized quintile bins (each containing *∼*277 coding sequences) and analyzed SD-like sequence conservation separately within each set.

## Protein structural data

Protein domain annotations were downloaded from Ciryam *et al.* [41]. We cross referenced annotations between our dataset and theirs, and for each annotated domain analyzed SD-like sites that occurred within 150 nucleotides downstream of the domain end (while maintaining previous restrictions on 5ʹ and 3ʹ gene ends). Control sites were selected from anywhere within the same gene (outside of SD-like sequences).

## Acknowledgments

The authors wish to thank Stephanie J. Spielman for help with nucleotide rate calculations using HyPhy. This work was supported by NIH grant R01 GM088344, NSF (Cooperative agreement no. DBI-0939454, BEACON Center), and Army Research Office (ARO, http://www.arl.army.mil/) grant W911NF-12-1-0390. LANA and MCJ acknowledge a gift from Leslie and John McQuown. MCJ acknowledges further support from the David and Lucile Packard Foundation, and the Camille-Dreyfus Teacher-Scholar Program.

## Supplementary Information: Selection removes Shine-Dalgarno-like sequences from within protein coding genes

**Supplementary Table S1:** SD sequence motif occurrence throughout the *E. coli* genome. Data shown for annotated protein coding genes that: i) have a nucleotide length is a multiple of 3, ii) are between 60 and 1000 amino acids in length, iii) contain no internal stop codons, and iv) end with a canonical stop codon. Genes with an upstream SD sequence were defined according to the presence of a 6-nt motif between -20 to -4 (relative to the start codon) that pair with the aSD sequence (5ʹ-CCUCCU-3ʹ) with a binding free energy less than the indicated threshold. Expectation was determined by shuffling the nucleotides between -20 to -1 for each gene within a genome, calculating as above, and repeating 100 times. SD-like sequences were similarly defined after excluding the first and last 60 nucleotides of each gene, testing each sequential 6 nucleotide motif, and counting the number of strong binding sequences. We only considered SD-like sequences where the binding energy was less than the defined threshold *and* was less than the two immediate neighboring sequences (i.e. motifs that are shifted one nucleotide up and downstream) to avoid double counting strong SD sequences that may have a signal in multiple sequential motifs. Expectation for SD-like sequences was performed by shuffling synonymous codons within each gene (preserving amino acid sequences, GC content, and gene-speciifc codon usage biases), calculating the number of SD-like sites for one instance of this shuffled genome, and repeating this procedure 100 times.

|  | Observed | Expected | *p*-value |
| --- | --- | --- | --- |
| Total protein coding genes | 4127 | - | - |
| Threshold (-4.5 kcal/mol) |  |  |  |
| Genes with upstream SD sequence | 1998 | 638.57 | $< 10^{-16}$ |
| SD-like sites within protein coding genes | 25001 | 30397.57 | $< 10^{-16}$ |
| Threshold (-3.5 kcal/mol) |  |  |  |
| Genes with upstream SD sequence | 2806 | 1129.72 | $< 10^{-16}$ |
| SD-like sites within protein coding genes | 55242 | 57802.7 | $< 10^{-16}$ |
| Threshold (-5.5 kcal/mol) |  |  |  |
| Genes with upstream SD sequence | 1502 | 429.05 | $< 10^{-16}$ |
| SD-like sites within protein coding genes | 11355 | 15864.69 | $< 10^{-16}$ |

**Supplementary Table S2:**
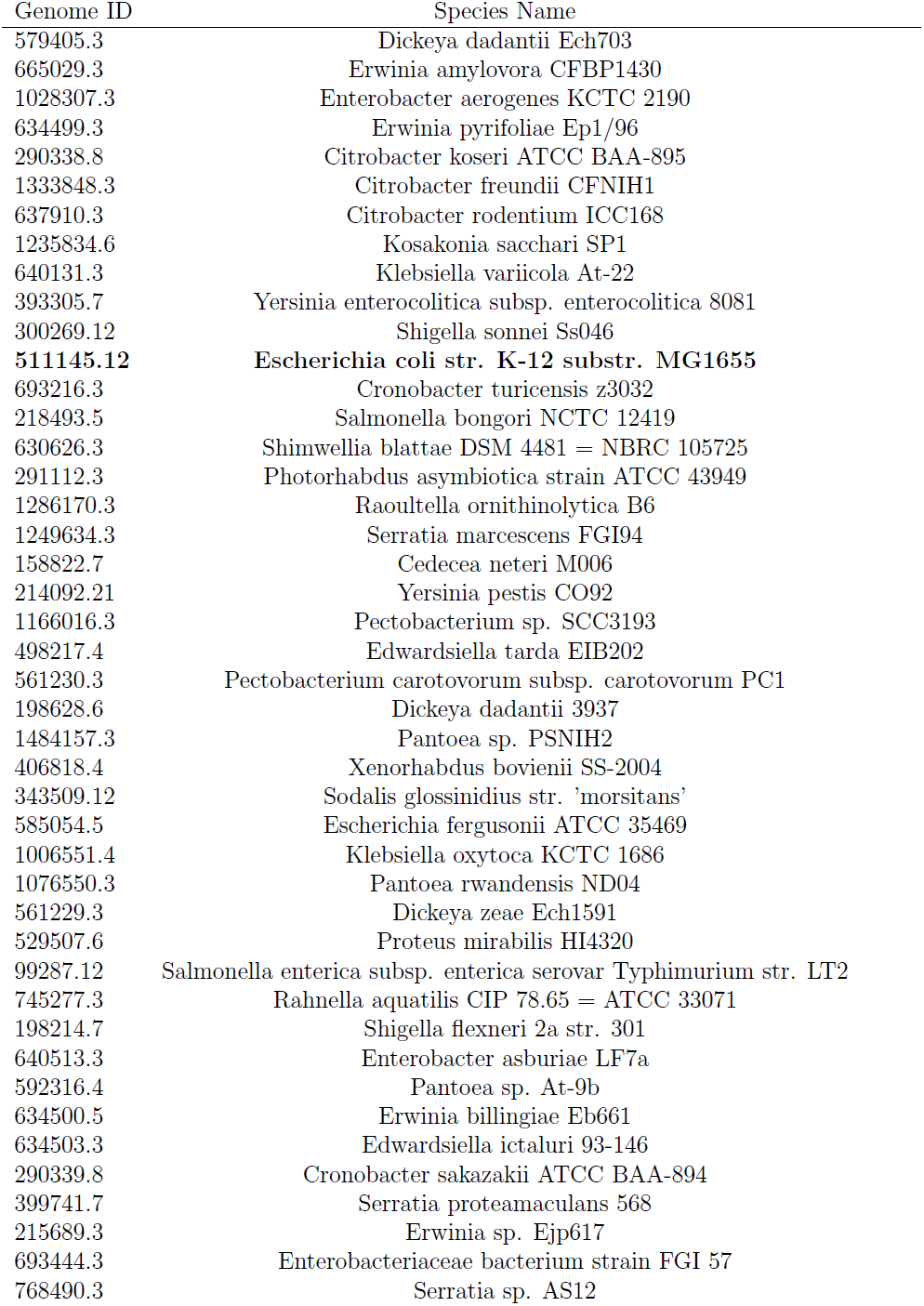
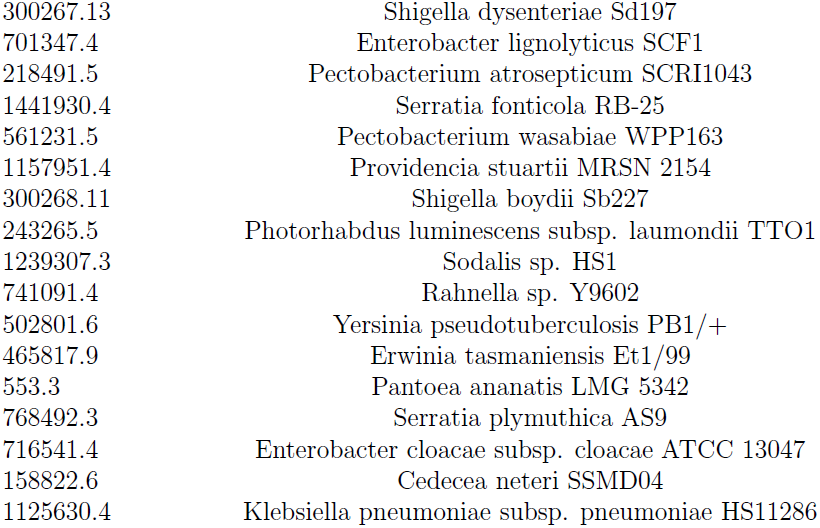
List of genomes analyzed in this study.

**Supplementary Figure S1:**
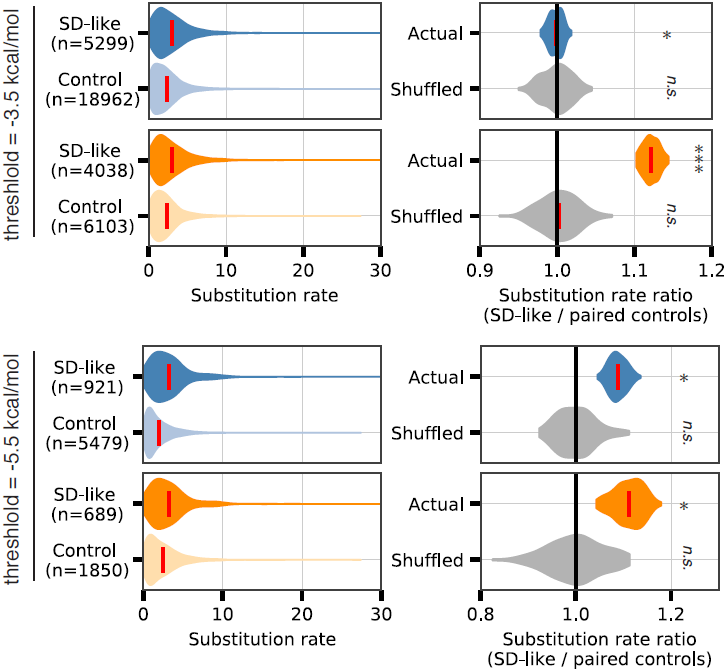
As in Fig. 1B, C. Here we show similar results for weaker (top) and stronger (bottom) thresholds for defining SD-like sequences. All patterns remain similar to those depicted in Fig. 1 with the exception of synonymous codon controls for the weakest thresholds, which show no substantial difference in substitution rate patterns and only borderline significance.

**Supplementary Figure S2:**
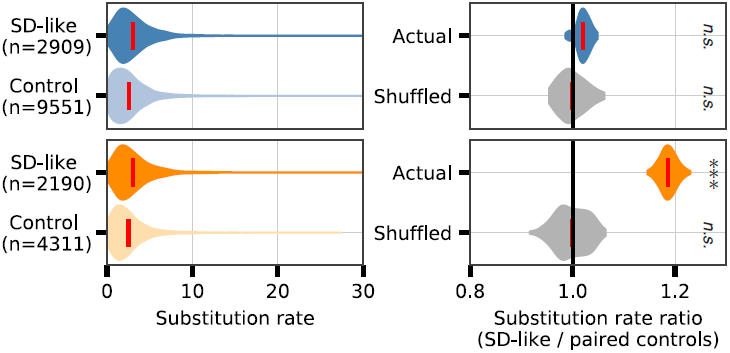
As in Fig. 1B,C. Here we show similar results when using *Y. pestis* as a reference genome to determine the location of SD-like sites. The exception being that, though slightly elevated, there is no significant difference in substitution rates between SD-like sites according to the synonymous codon control. (***denotes *p <* 0.001)

**Supplementary Figure S3:**
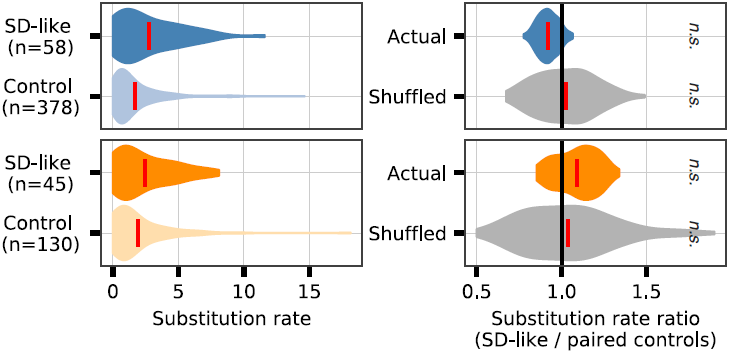
Using the same methodology as in Fig. 1D,E. Here we analyzed putative SD sites in the 3ʹ terminus of genes that are *not* directly followed by an annotated coding sequence. Due to the fact that these sequences are likely not acting as true SD sequences despite being in the 3ʹ terminus, we expected and observed no significant difference in substitution rates according to either null model.

**Supplementary Figure S4:**
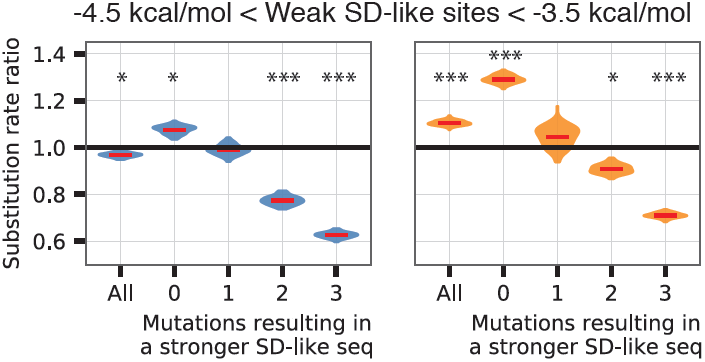
Weak SD-like sites follow the same patterns of substitution rate ratios as SD-like sites. All sites analyzed here are weaker in absolute terms compared to the weakest sites analyzed in Fig. 2. Nevertheless, we still observe elevated substitution rate ratios in the sites that are *locally* strong compared to those that are *locally* weak. In particular, locally strong sites here (where no mutations will result in a stronger SD-like sequence) exceed the substitution rate ratios of sites depicted in Fig. 2 that are stronger in the absolute sense (more negative ∆*G*) but *relatively* weak given their local mutational context (any mutation will result in stronger SD-like sequence). (* denotes *p <* 0.05, *** denotes *p <* 0.001)

**Supplementary Figure S5:**
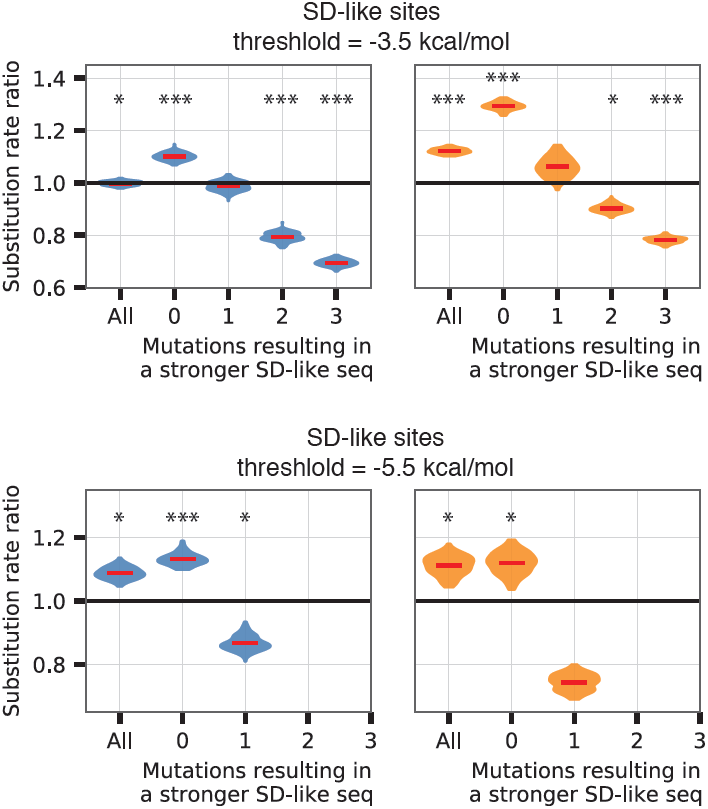
As in Fig. 2. Here we show similar results for weaker (top) and stronger (bottom) thresholds for defining SD-like sequences. Breaking mutations up by their local mutational context reveals that the patterns for individual classes remain un-changed from conclusions presented in the main text. For the most stringent SD-like sequence threshold (bottom), no locally-weak sites are strong enough to be analyzed, thus categories “2” and “3” are empty. (* denotes *p <* 0.05, *** denotes *p <* 0.001)

**Supplementary Figure S6:**
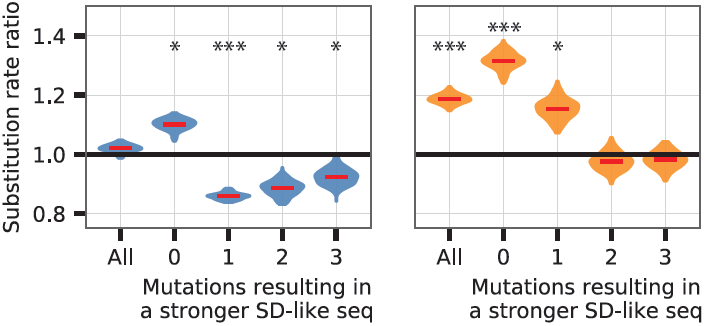
As in Fig. 2. Here we show similar results when using *Y. pestis* as the reference genome to determine the location of SD-like sites. (* denotes *p <* 0.05, *** denotes *p <* 0.001)

**Supplementary Figure S7:**
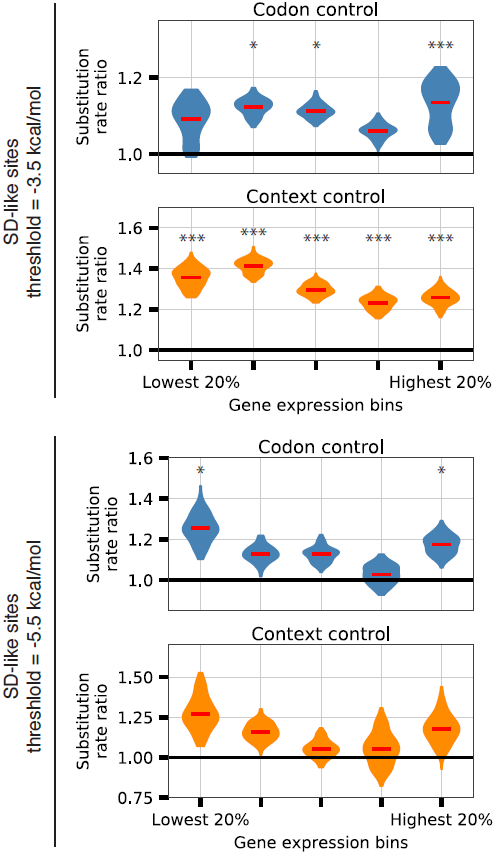
As in Fig. 3. Here we show similar results for weaker (top) and stronger (bottom) thresholds for defining SD-like sequences. For stringent thresholds, very few analyzable sites remain within each bin and statistical significance is frequently not observed.

**Supplementary Figure S8:**
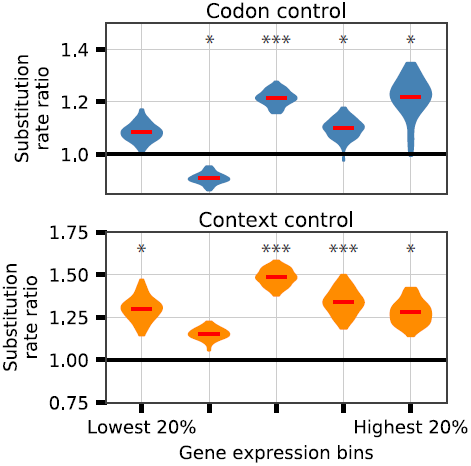
As in Fig. 3. Here we show similar results when using *Y. pestis* as the reference genome to determine the location of locally strong SD-like sites. Note that protein homologs are still partitioned into bins according to their measured abundances in *E. coli*. (* denotes *p <* 0.05, *** denotes *p <* 0.001)

**Supplementary Figure S9:**
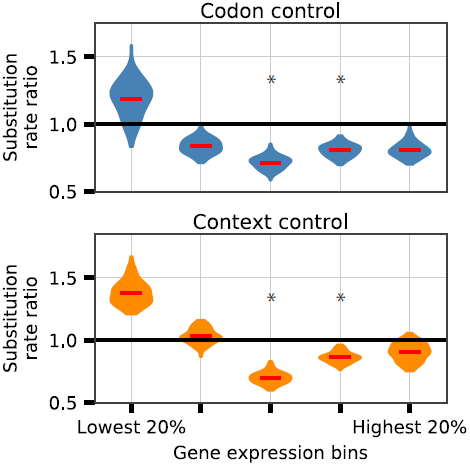
As in Fig. 3, regardless of protein abundance bin, substitution rate ratios for *locally weak sites* are inconsistent with the hypothesis that the highest abundance proteins may contain more evolutionarily constrained SD-like sequences. (^*^ denotes *p <* 0.05)

**Supplementary Figure S10:**
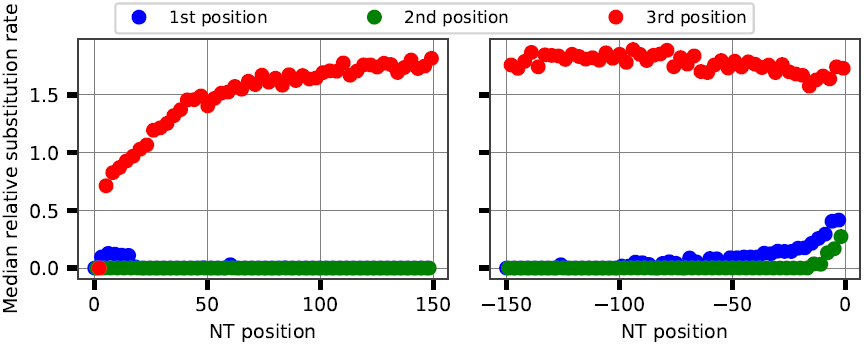
Substitution rates using a meta-gene analysis follow expected patterns of sequence conservation. Notably, 3rd position nucleotides have substantially elevated median substitution rates and substitution rates are lower towards the 5ʹ end of coding sequences.

**Supplementary Figure S11:**
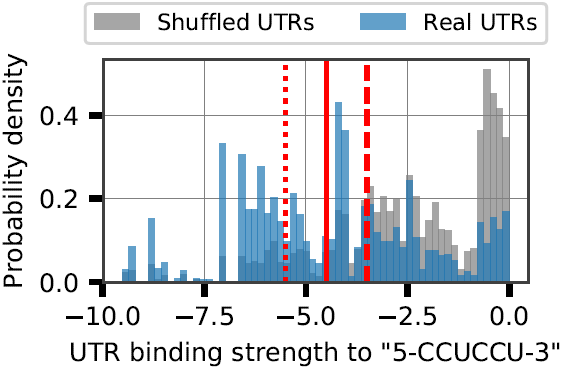
Distribution of the binding strengths for the strongest aSD sequence binding hexamer between positions -20 to -1 for each protein coding gene in *E. coli* (blue). Shown in grey is the expected distribution of binding strengths when first shuffling the 20 nucleotides upstream of each gene. The solid red line depicts the threshold used throughout the main text to determine SD-like sequences (-4.5 kcal/mol), while dotted and dashed lines to the left and right depict more and less stringent thresholds used to test the robustness of findings (-5.5 and -3.5 kcal/mol).

